# Conserved recombination patterns across coronavirus subgenera

**DOI:** 10.1101/2021.11.21.469423

**Authors:** Arné de Klerk, Phillip Swanepoel, Rentia Lourens, Mpumelelo Zondo, Isaac Abodunran, Spyros Lytras, Oscar A MacLean, David Robertson, Sergei L Kosakovsky Pond, Jordan D Zehr, Venkatesh Kumar, Michael J. Stanhope, Gordon Harkins, Ben Murrell, Darren P Martin

**Author notes:** Address for correspondence: Arné de Klerk, Division Of Computational Biology, Faculty of Health Sciences, University of Cape Town, Anzio Road, 7925 Observatory, Cape Town, South Africa.

## Abstract

Recombination contributes to the genetic diversity found in coronaviruses and is known to be a prominent mechanism whereby they evolve. It is apparent, both from controlled experiments and in genome sequences sampled from nature, that patterns of recombination in coronaviruses are non-random and that this is likely attributable to a combination of sequence features that favour the occurrence of recombination breakpoints at specific genomic sites, and selection disfavouring the survival of recombinants within which favourable intra-genome interactions have been disrupted. Here we leverage available whole-genome sequence data for six coronavirus subgenera to identify specific patterns of recombination that are conserved between multiple subgenera and then identify the likely factors that underlie these conserved patterns. Specifically, we confirm the non-randomness of recombination breakpoints across all six tested coronavirus subgenera, locate conserved recombination hot- and cold-spots, and determine that the locations of transcriptional regulatory sequences are likely major determinants of conserved recombination breakpoint hot-spot locations. We find that while the locations of recombination breakpoints are not uniformly associated with degrees of nucleotide sequence conservation, they display significant tendencies in multiple coronavirus subgenera to occur in low guanine-cytosine content genome regions, in non-coding regions, at the edges of genes, and at sites within the Spike gene that are predicted to be minimally disruptive of Spike protein folding. While it is apparent that sequence features such as transcriptional regulatory sequences are likely major determinants of where the template-switching events that yield recombination breakpoints most commonly occur, it is evident that selection against misfolded recombinant proteins also strongly impacts observable recombination breakpoint distributions in coronavirus genomes sampled from nature.

## Introduction

Coronaviruses are a family of vertebrate-infecting single-stranded, positive-sense RNA viruses with genomes ^~^27-32kb in length. The family has four genera - *Alphacoronavirus, Betacoronavirus, Gammacoronavirus* and *Deltacoronavirus* - each of which has been further subdivided into a number of subgenera such as *Pedacovirus* in the genus *Alphacoronavirus*, and *Merbecovirus, Embecovirus, Nobecovirus*, and *Sarbecovirus* in the genus *Betacoronavirus*, and *Igacovirus* in the genus *Gammacoronavirus* (Coronaviridae - Positive Sense RNA Viruses - Positive Sense RNA Viruses (2011) -ICTV 2011). Besides SARS-CoV-2, the *Sarbecovirus* member that causes COVID-19, there are four other known *Betacoronavirus* lineages and two known *Alphacoronavirus* lineages that either cause - or have caused - epidemiologically significant disease outbreaks in humans.

Coronavirus genomes generally contain seven to ten genes, often with varying arrangements and compositions (Fig. 1) (Lai 1996). The largest gene, ORF1ab, encodes multiple non-structural proteins which are involved in viral transcription, replication, proteolytic processing, modulation of host gene expression and the suppression of host immune responses (Emam *et al*. 2021). Directly downstream of ORF1ab in most known coronavirus genomes is the Spike (S) gene (although in *Embecoviruse*s, for example, a haemagglutinin-esterase gene separates ORF1ab and the S gene). Spike is the structural glycoprotein found on the outside of coronavirus particles that gives them their iconic crown-like protrusions. Spike binds to cell membrane receptors and mediates virus entry into host cells. It is considered a class I fusion protein in that it contains both a receptor-binding domain (called S1) and a domain for mediating the membrane fusion process (called S2) (White *et al*. 2008; Xia *et al*. 2020). The Spike of SARS-CoV-2 has become a target in the development of vaccines and therapeutic drugs for COVID-19, due both to its importance during the viral infection cycle and to its being the primary target of host immune responses (Krumm *et al*. 2021).

**Figure 1.**
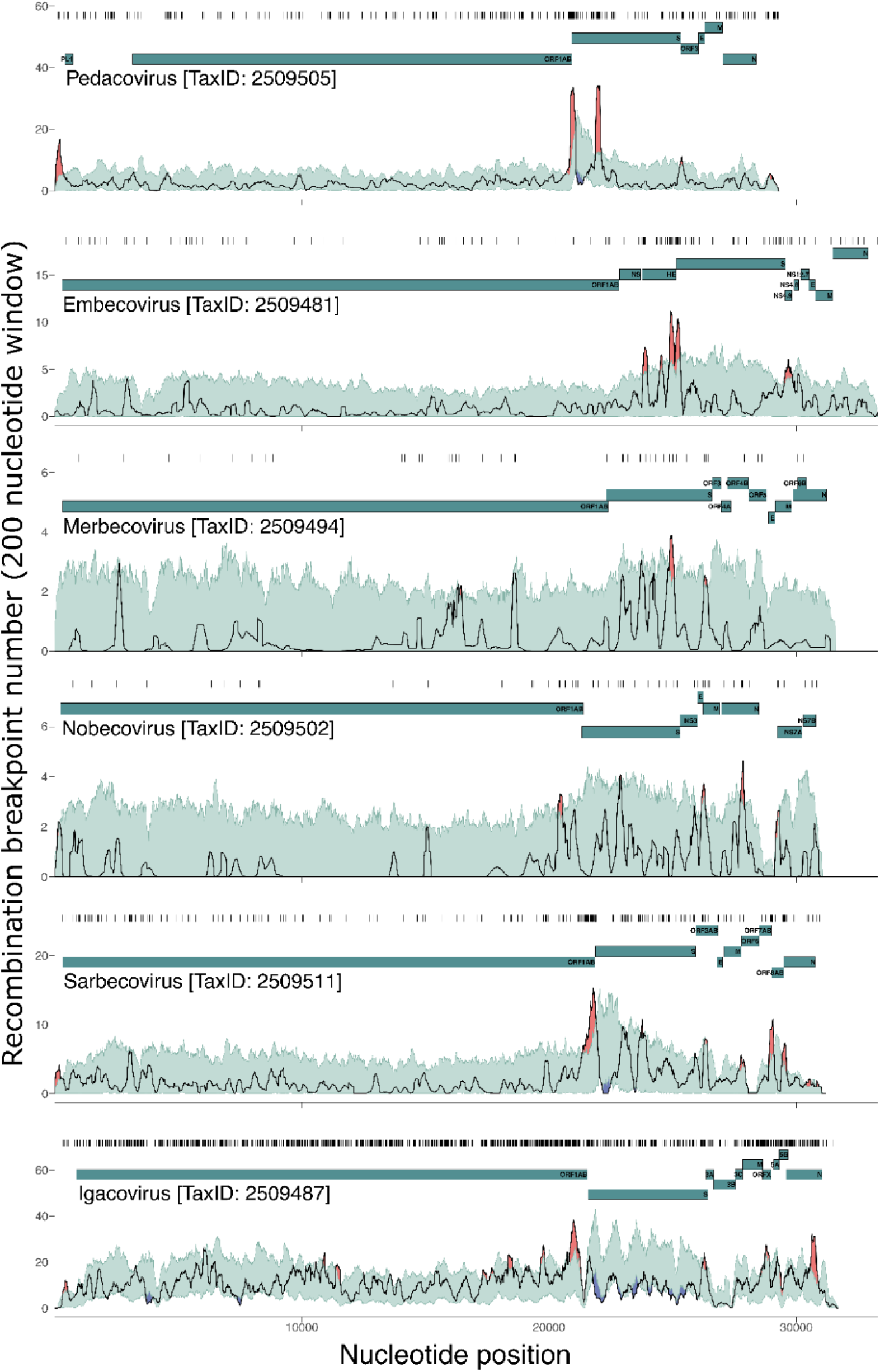
Variation across coronavirus genomes in the densities of detectable recombination breakpoints. All detected breakpoint positions are indicated directly above each graph with black vertical lines. A gene-map is shown as teal-coloured lines. The light green areas indicate 99% bounds of expected degrees of breakpoint clustering under random recombination. Areas where the black lines (breakpoint numbers per 200 nucleotide window) have emerged above the green areas, are considered potential recombination hot-spots, and are marked in red. Areas where the black lines drop below the green areas are considered potential recombination cold-spots, and are marked in blue.

Although coronaviruses have low mutation rates relative to those of other single-stranded RNA viruses (Denison *et al*. 2011; Jaroszewski *et al*. 2021), coronavirus populations are characterized by high degrees of genetic diversity (Liu *et al*. 2017). Much of this genetic diversity is likely generated and maintained by high rates of within-species (Dudas and Rambaut 2016; Su *et al*. 2016; Anthony *et al*. 2017; Forni, Cagliani and Sironi 2020) and between-species genetic recombination (Wesley 1999;Decaro *et al*. 2015; Wang *et al*. 2015, 2020). The first credible reports of recombination in coronaviruses were made in the mid-to-late 1980s and focused on mixed *in vitro* and *in vivo* infections of different Murine mouse hepatitis virus strains (Makino *et al*. 1986; Keck *et al*. 1988a, 1988b; Banner and Lai 1991). By the year 2000, comparative analyses of coronavirus genomes sampled from natural infections had yielded convincing evidence that recombination, particularly between divergent coronaviruses within individual subgenera, is a major contributor to coronavirus evolution (Kusters *et al*. 1990; Wang, Junker and Collisson 1993; Jia *et al*. 1995; Lee and Jackwood 2000). For example, a complex recombinant history is evident between the Alphacoronavirus-1 species involving canine coronavirus, that primarily infects dogs (however, see recent exceptions of canine CoV infections in human: (Lednicky *et al*. 2021; Vlasova *et al*. 2021; Zehr *et al*. 2021)), transmissible gastroenteritis virus, infecting pigs and feline coronavirus from cats (Herrewegh *et al*. 1998; Decaro *et al*. 2009).

The most common mechanism of recombination in coronaviruses (and many other RNA viruses too) is known as copy-choice, where a viral RNA dependent RNA polymerase (RdRp) is interrupted during replication, drops off the RNA template that it was copying, and re-engages with a different RNA template at a homologous position before resuming replication (Cheng and Nagy 2003). Such template switches during replication yield recombinant daughter genomes with different regions of sequence being derived from two different “parental” genomes. The genome sites at which template switches occur are referred to as recombination breakpoints.

Recombination likely provides coronaviruses with more evolutionary options than would be available to them by mutation alone (Crameri *et al*. 1998; Simon-Loriere *et al*. 2009). While it is expected that many newly arising mutations within genetically compact viral genomes (such as those of coronaviruses) will have negative fitness consequences, so too will many of the recombination events that occur between genetically divergent genomes (Drummond *et al*. 2005). By transferring pieces of genomes into genomic backgrounds with which they did not coevolve, recombination will frequently run the risk of disrupting favourable coevolved intra-genome interactions (commonly referred to as epistatic interactions). Examples of favourable coevolved intra-genome interactions that could be disrupted by recombination include those between nucleotides that base-pair to form biologically functional genomic secondary structures, those between pairs of amino acids that interact to mediate protein folding, those between the binding domains on protein surfaces that mediate multi-protein complex formation, and those between sequence-specific nucleic acid binding domains and nucleotide sequence motifs that mediate gene regulation and genome replication (Martin *et al*. 2005b). However, since recombination generally occurs between fully functioning genomes, the range of potential negative fitness consequences of recombination are, in general, expected to be less extreme than those that might occur due to newly arising mutations (Drummond *et al*. 2005). In fact, genetic recombination between closely related viruses almost certainly helps defend against the accumulation within genomes of mildly deleterious mutations that, in high enough numbers, might otherwise have serious fitness consequences (Muller 1964; Woo *et al*. 2010;Hussin *et al*. 2015; Goldstein *et al*. 2021).

Here we analyse patterns of recombination evident in whole-genome datasets drawn from one *Alphacoronavirus* subgenus, one *Gammacoronavirus* subgenus and four *Betacoronavirus* subgenera. We confirm previous reports that natural recombination between genetically divergent coronaviruses is common and find strong evidence that detectable recombination breakpoint sites are not randomly distributed across coronavirus genomes. Specifically, we demonstrate the likely occurrence of breakpoint hot- and cold-spots, some of which are conserved across multiple coronavirus groups. Further, we find detectable associations across multiple different coronavirus subgenera between recombination breakpoint locations and various sequence features that might impact the mechanistic predisposition of certain genome sites to recombine more than others (such as decreased guanine-cytosine content, and the locations of transcriptional regulatory sequences). We also find evidence across multiple subgenera that selection differentially favours the survival of recombinants based on the genome sites at which breakpoints occur (such as at the edges of genes or in intergenic regions relative to the middle portions of genes).

## Methods

### Data collection

All publicly available near full-length genomic sequences for viruses in six well-sampled coronavirus subgenera (*Igacovirus, Embecovirus, Merbecovirus, Nobecovirus, Pedacovirus* and *Sarbecovirus*) were downloaded from the NCBI Virus (Hatcher *et al*. 2017), CNCB (Song *et al*. 2021), and CoVDB (Zhu *et al*. 2021) databases between February and May of 2021. Each of the six subgenus-level datasets was aligned with MAFFT using default settings (Katoh and Standley 2013). All but one sequence in groups of sequences sharing more than 99% nucleotide sequence identity were removed to yield datasets for recombination analysis containing between 16 and 412 genome sequences sharing ≥75% similarity (Supplementary Table 1; Supplementary data).

### Recombination detection

Recombination was detected and analysed using RDP5 (Martin *et al*. 2021) with default settings except that sequences were treated as linear. Each of the six coronavirus datasets were analysed for recombination using a fully exploratory automated scan with the RDP (Martin and Rybicki 2000), GENECONV (Sawyer 1989), and MaxChi (Maynard Smith 1992) methods to detect recombination signals (i.e these were used as “primary scanning methods”), and the Bootscan (Martin *et al*. 2005a), Chimaera (Pettersen *et al*. 2004), SiScan (Gibbs, Armstrong and Gibbs 2000) and 3Seq (Lam, Ratmann and Boni 2018) methods to verify the signals (these latter four methods being used as “secondary scanning methods”). From among the individual recombination signals that were each detectable by four or more of these methods, RDP5 refined the positions of detected recombination breakpoints using a hidden Markov model (HMM) based approach (described in detail in the RDP manual at http://web.cbio.uct.ac.za/~darren/RDP4Manual.pdf) and determined a plausible near minimal subset of unique recombination events that would be needed to account for all of the detected recombination signals. Each of the unique recombination events detected by RDP5 in each of the six analysed coronavirus subgenera datasets was characterized by: (1) a 5’ and 3’ pair of maximum likelihood breakpoint locations and their associated probability distributions, (2) a list of one or more sequences carrying evidence of the recombination event (multiple sequences can have evidence of the same recombination event if the event occurred in a common ancestor), and (3) a list of analysed sequences that are closely related enough to the actual parents of the recombinant that they could be used as proxies for the actual parents to detect the recombination events. The overall-recombination patterns in the six subgenera datasets were visualized using recombination region count matrices produced using RDP5. These matrices indicate the numbers of detected recombination events that separated individual genome sites from all other genome sites.

### Recombination breakpoint hot- and cold-spot tests

For each of the subgenera, a recombination breakpoint distribution map was constructed from the lists of 5’ and 3’ breakpoint probability distributions associated with each detected recombination event. This was done by sliding a 200 nt window, one nucleotide at a time, along the full length of the analysed alignment, summing the probabilities of all identified breakpoints falling within the window, and plotting these counts at the nucleotide coordinate at the centre of the window. A permutation test implemented in RDP5, described previously (Heath *et al*. 2006; Lytras *et al*. 2021), was then used to identify recombination breakpoint clustering patterns that varied significantly from expectations under random recombination. Briefly, this test involved: (1) randomly shuffling the breakpoint locations of each of the observed recombination events to maintain the spacing between 5’ and 3’ breakpoint pairs with respect to the numbers of polymorphic nucleotide sites separating the breakpoint pairs within the triplets of analysed sequences, used to detect the recombination event; (2) ensuring that in instances where individual sequences contained evidence of multiple independent recombination events, the regions bounded by 5’ and 3’ breakpoint pairs for those events did not overlap to a greater or lesser degree those observed in the actual recombinants (i.e. the spacings of all the 5’ and 3’ breakpoint locations of all overlapping events within a single sequence were maintained); (3) ensuring that in instances where breakpoints were flagged as having undetermined positions in the actual dataset (such as breakpoints called at the start/end of the alignment or at sites that were overprinted by subsequent recombination events), these were excluded from breakpoint counts; (4) making recombination breakpoint distribution maps for each permuted dataset using the exact same approach as that used for the actual dataset; and (5) identifying unusually high or low degrees of breakpoint clustering in the actual dataset as those window coordinates where the breakpoint probability sums of the actual dataset fell outside the bounds of those determined at that coordinate for 99% of the permuted datasets. With this test, unusually high degrees of breakpoint clustering (i.e. greater than 99% of the permuted datasets at a given genome site) are suggestive of recombination hot-spots, whereas unusually low degrees of clustering (i.e. lower than 99% of the permuted datasets at a given genome site) are suggestive of recombination cold-spots.

It is important to stress, that this breakpoint clustering test is not conservative; given the lengths and degrees of diversity of the analysed coronavirus genomes, it is expected that one or two hot-spot-like clusters of breakpoints would be detectable in each of the datasets even under completely random recombination (Lytras *et al*. 2021). We therefore referred to hot-spots detected by this test in individual datasets as “potential hot-spots” and required that for a particular genome site to be defined as an actual statistically-supported hot-spot, potential hot-spots needed to be detectable at a homologous site in two or more of the different analysed subgenus datasets.

### Comparing recombination breakpoint counts between pairs of pre-defined genome regions

We used a version of the breakpoint clustering hot- and cold-spot test that compared observed breakpoint numbers in two preselected groups of sites in an analysed alignment (Lefeuvre *et al*. 2009). Since the original recombination breakpoint distribution test determined whether the numbers of breakpoints observed in 200 nt sliding windows were greater or lesser than chance under random recombination, the test relied on the detection of sufficient breakpoints for statistically implausible clusters of breakpoints to emerge. As the number of detected recombination breakpoints varied widely between the different coronavirus datasets (ranging from 65 for the *Merbecoviruses* and 1703 for the *Igacoviruses*), the power of the test varied substantially. In an adapted version of the test, we partitioned the sites in each of the six datasets into two large subsets and directly compared observed breakpoint numbers in each of the site subsets to those expected under random recombination. We specifically compared densities of breakpoints falling at: (1) non-protein-coding sites vs protein-coding sites; (2) the beginning and ending 5% of sites within individual protein-encoding regions vs the middle 90% of these regions (in the case of ORF1ab we defined protein encoding-regions as those encoding individual post-translational protein cleavage products), (3) genome sites encoding a particular protein vs those encoding all other proteins within the genome and (4) sites within a specified number of nucleotides (2, 9, 21, 46) of a transcriptional regulatory sequence vs those in the remainder of the genome.

### Testing for associations between GC content or pairwise sequence similarity and recombination breakpoint sites

A further modification of the breakpoint clustering hot- and cold-spot test was used to test for associations between breakpoint sites (specifically breakpoint probability distributions) and: (1) guanine+cytosine (GC) content; and (2) pairwise sequence similarity (Simon-Loriere *et al*. 2010). In this test average GC proportions or pairwise sequence similarities of sites between a specified number of nucleotides (either 10 or 20) of every site in the genome across all possible sequence pairs were determined. Breakpoint probabilities at each site were multiplied with the GC proportion or pairwise similarity associated with that site and summed across all sites. These sums for the real datasets were then compared with the corresponding sums from the permuted datasets. For each analysed subgenus dataset the proportion of permuted datasets with sums higher than or equal to the real dataset were reported as the probability that there was no association between breakpoint positions and either higher GC proportions or higher degrees of pairwise similarity. Conversely, the proportion of permuted datasets with sums lower than or equal to those determined for the real dataset was reported as the probability that there was no association between breakpoint positions and either lower GC proportions or lower degrees of pairwise sequence similarity.

### Identification of potential transcriptional regulatory sequences

SuPER was used to detect transcriptional regulatory sequence leader (TRS-L) sites and a custom Python (Rossum and Drake 2010) script was used to infer transcriptional regulatory sequence body (TRS-B) sites (Yang *et al*. 2021). For the algorithm implemented in SuPER to infer the subgenomic mRNA positions without RNA-seq data, annotation files and reference sequence files were downloaded from NCBI in September 2021 for the best-sampled species in each of the six coronavirus subgenus datasets. A Python script (https://github.com/phillipswanepoel/trsb-finder) was used to search for potential TRS-B sites in each subgenus dataset, following the methodology used in SuPER, which involved searching for all occurrences of sub-sequences with a Levenshtein distance of one or zero from the TRS-Leader sequence (Yang *et al*. 2021). These potential TRS-B sites were then filtered, removing all the sites not conserved across at least 75% of the analysed sequences and removing upstream sites when multiple sites were found in close proximity 5’ of the start of the same ORF. This filtered siteset was then tested for association with breakpoint positions in each of the six analysed subgenera datasets using RDP5.

Given that neither the TRS distributions nor the breakpoint distributions were random in any of the analysed datasets and that both TRS sites and recombination breakpoint clusters occurred at the edges of coronavirus genes, we anticipated that the association test could have a high false-positive rate. To estimate the false discovery rate (FDR) of the breakpoint association test, a custom Python script (https://github.com/phillipswanepoel/trsb-finder) was used to generate randomly permuted versions of TRS-B site locations, for each of the subgenera alignments. As input, the script takes a coronavirus subgenus alignment and associated TRS-B sites, then outputs an RDP5 readable siteset file containing the permuted nucleotide positions. These positions are calculated by collectively shifting all the TRS-B sites by some number of nucleotides (which preserves their spacing), varied randomly between one and the length of the analysed alignment. If a new shifted TRS position was beyond the end of the genome, the position was “wrapped” around to the other end of the genome. Two hundred permuted TRS-B site-sets were tested and the average estimated FDR across all datasets for the association between breakpoint and TRS sites was 17.27% (i.e. 17.27% of the analyses with “shifted” TRS-B sites yielded a significant association - with a p-value < 0.05 - between these sites and observed breakpoint positions). Given that the FDRs for individual subgenera datasets ranged from 5% to 29%, we only considered associations detected between TRS-B and breakpoint sites as being significant if they were detected in multiple datasets.

### Protein folding disruption test

To test whether the observed recombination events were less disruptive of protein folding than would be expected if recombination breakpoints were randomly distributed, the SCHEMA test (Meyer *et al*. 2003; Lefeuvre *et al*. 2007), implemented in RDP5, was used to examine all protein-coding regions with associated publicly available high resolution atomic coordinate data (obtained from the Protein Data Bank; https://www.rcsb.org/ (Berman, Henrick and Nakamura 2003)) and within which more than ten recombination breakpoints were detected. These stipulations were required to ensure that the test would have sufficient power to detect whether observed recombinants displayed significantly lower degrees of protein folding disruption with the SCHEMA test than would be expected under random recombination. Of all 56 unique encoded proteins for which structural data was available (across all subgenera), only Spike was amenable to further analysis. Specifically, four subgenera (*Pedacovirus, Merbecovirus, Sarbecovirus* and *Igacovirus*) had both available Spike atomic coordinate structural data and >10 detected recombination breakpoints in the portion of the S gene corresponding to the structural data.

The SCHEMA test involves identifying potential interactions that occur between amino acid residues within folded proteins (in our case pairs of non-hydrogen atoms from different amino acids within 4.5 Å of one another) and counting the numbers of interacting amino acid pairs within a chimaera of two parental amino acid sequences, where the chimaera has a different pair of amino acids than both parents. The 4.5 Å interaction cut-off (the default setting) corresponds to approximately five to eight potential pairwise interactions per residue. The counts of potentially altered pairwise amino acid interactions (called the disruption or E-score) that the SCHEMA method calculates has been shown to strongly correlate with observed degrees of fold disruption within chimaeric proteins (Meyer *et al*. 2003). To determine whether observed recombinants expressed chimeric proteins with significantly lower E-scores than expected under random recombination, we used the permutation-based recombinant protein simulation approach of Lefeuvre et al. (2009).

## Results and Discussion

### Conserved recombination breakpoint hot- and cold-spots within coronavirus genomes

Using a combination of recombination detection methods implemented in RDP5, we identified 416 unique recombination events in the *Pedacovirus* dataset, 255 in *Embecovirus*, 65 in *Merbecovirus*,107 in *Nobecovirus*, 282 in *Sarbecovirus*, and 1703 in *Igacovirus*. The variable numbers of detected recombination events between datasets should not be considered evidence that the viruses in some subgenera recombine more than others. Rather, the variable numbers reflect differences in both the numbers of analysed sequences in each dataset (e.g. the *Igacovirus* dataset had the most sequences) and the genetic diversity of the sequences in the different datasets (e.g. the *Pedacovirus* dataset had the least diverse sequences; Supplementary Table 1).

To visualise the recombination breakpoints associated with these events in each subgenus, breakpoint distribution plots (Figure 1), and recombination region count matrices (Figure 2) were constructed. The breakpoint distribution plots revealed clusters of breakpoints that were either more or less dense at individual genome sites than those observed at corresponding sites in 99% of permuted datasets where recombination breakpoint positions were randomly distributed (Figure 1). Potential recombination hot-spots were detected in all of the analysed subgenera (indicated by red shading in Figure 1) and recombination cold-spots in three of them (indicated by blue shading in Figure 1).

**Figure 2.**
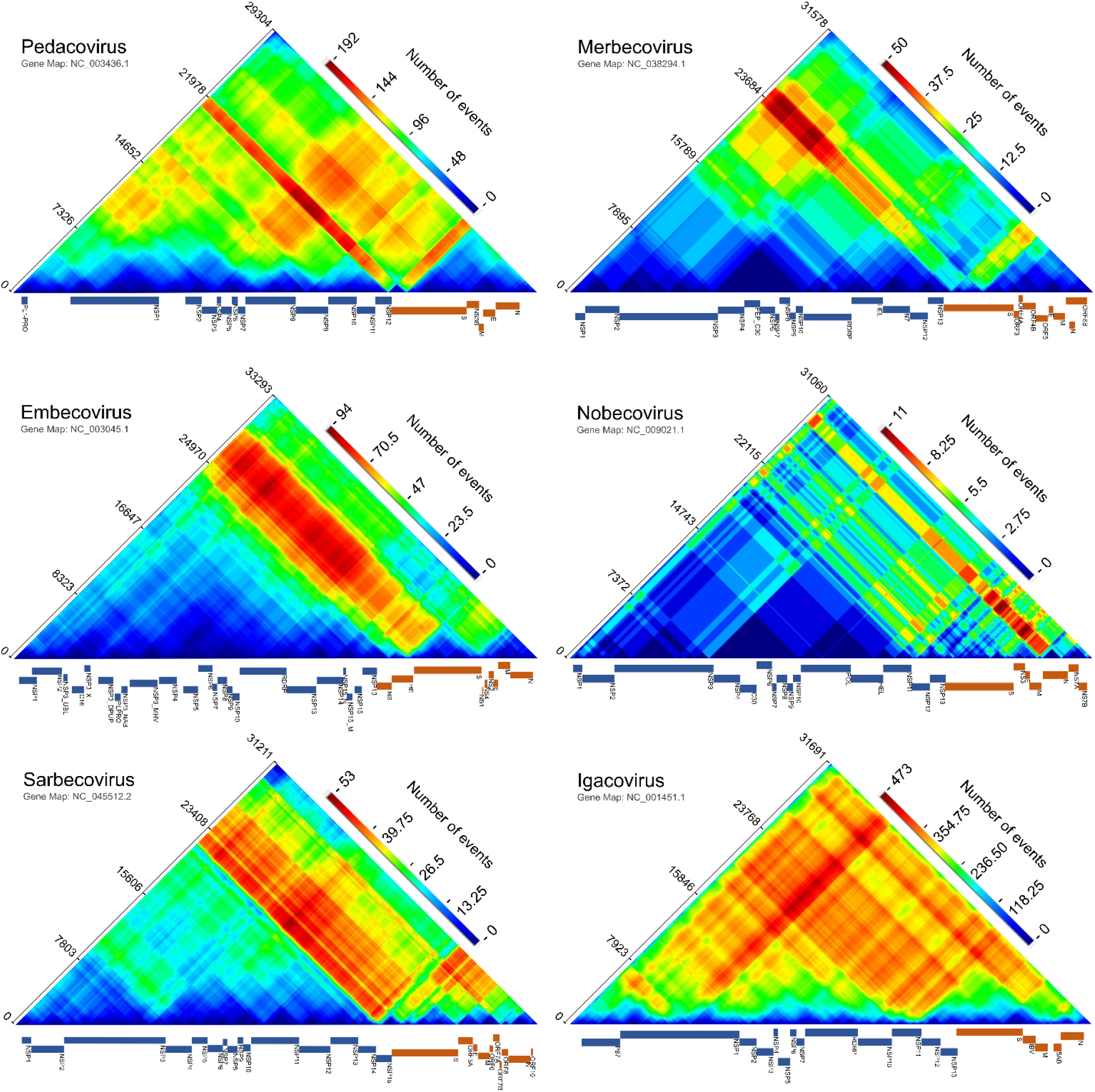
Recombination region count matrices indicating genome regions that are most and least commonly transferred during detectable coronavirus recombination events. Unique recombination events for six coronavirus subgenera, mapped onto recombination region count matrices based on determined breakpoint positions. Each cell in the matrix represents a pair of genome sites with the colours of cells indicating the numbers of times recombination events separated the represented pairs of sites. Reference sequence gene maps of the most prevalent virus in each subgenus were obtained from the NCBI nucleotide database (https://www.ncbi.nlm.nih.gov/nuccore) and are plotted alongside each matrix. Nucleotide positions are plotted according to full analysed nucleotide sequence alignments (Supplementary material). Genome maps indicate the coding regions of individual protein products. Non-structural proteins encoded by ORF1ab are indicated in blue and other genes are indicated in orange.

In all the subgenera other than *Embecovirus* and *Merbecovirus*, potential recombination hot-spots were detected within 300 nucleotides of the 5’ end of the genome. This non-coding region is upstream of ORF1ab, where the transcription and replication initiation sites are. These initiation sites prime the transcription of subgenomic mRNAs and contain extensive secondary structures that are partially conserved amongst the viruses belonging to a given coronavirus genus (reviewed in (Yang and Leibowitz 2015) (Siegfried *et al*. 2014; Manfredonia *et al*. 2020)). In various other viruses such as HIV, recombination breakpoints tend to colocalize with highly structured genome regions (Simon-Loriere *et al*. 2010) and it is therefore plausible that recombination hot-spots detected at the 5’ end of *Pedacovirus, Nobecovirus, Sarbecovirus* and *Igacovirus* genomes might also be attributable to secondary-structure induced template-switching during replication.

Multiple other potential recombination hot-spots were detected near the boundaries of various different genes in the 3’ genome regions of *Sarbecoviruses, Nobecoviruses*, and *Igacoviruses*. In *Sarbecoviruses*, four potential recombination hot-spots were detected in the 3’ genome regions, between the M gene and ORF6, between ORF7AB and ORF8AB, and between ORF8AB and N. In *Nobecoviruses*, potential recombination hot-spots were detected in the 3’ genome regions, between the E gene and the M gene, and near the centre of the N gene. In *Igacoviruses* there were several potential hot-spots in the 3’ genome region, the clearest of which fell towards the 3’ end of N.

### Breakpoint distributions in and around the S gene are consistent with recombination facilitating host adaptation and/or immune evasion

Most noteworthy of all the detected potential breakpoint hot-spots were those falling within 800 nucleotides upstream of the S gene start codon in all subgenera other than the *Merbecoviruses*. The conserved arrangement of recombination breakpoint clusters in relation to the S gene likely underlies the observation that, according to our analyses, the S gene has been frequently transferred in its entirety during recombination events in *Igacoviruses, Sarbecoviruses* and *Embecoviruses* (note red diagonals associated with the S genes of these subgenera in Figure 2). However, in all analysed coronavirus groups other than the *Embecoviruses* and *Igacoviruses*, potential recombination hot-spots were also detected within the 5’ half of the S gene (S1 domain), further suggesting that the 3’ half of the gene (S2 domain) is the portion that is most commonly transferred during recombination events as a complete module (note the red diagonals associated with the 3’ part of the S gene in Figure 2). The locations of the detected recombination hot-spots in, and immediately adjacent to, the S gene suggest that either the complete S gene or its 3’ half, has been frequently transferred during recombination. This is not to say, that exceptions to this undoubtedly occur, including for example a likely instance in Alphacoronavirus-1 (a group not included in this current analysis, because of lower complete genome numbers) involving a section of the S2 domain of the newly described canine coronavirus HuPn-2018 (Zehr *et al*. 2021).

The Spike proteins that are encoded by the S gene are composed of an amino-terminal subunit 1 (S1) and a carboxyl-terminal subunit 2 (S2) (Wrapp *et al*. 2020). The S1 contains the N-terminal domain (NTD) and a receptor-binding domain (RBD) which mediates the binding of viral particles to host cell surface receptors. Different coronaviruses bind to different receptors. For example, the *Merbecovirus*, MERS-CoV, binds dipeptidyl peptidase-4 (DPP4), the *Pedacovirus*, PEDV, binds aminopeptidase N, and the *Sarbecoviruses* SARS-CoV and SARS-CoV-2 bind to angiotensin-converting enzyme 2 (ACE2) (Yeager *et al*. 1992; Li, Ge and Li 2007; Belouzard *et al*. 2012; Raj *et al*. 2013;Reusken *et al*. 2016; Wan *et al*. 2020). It is also likely that in some coronaviruses the NTD of Spike also interacts with cell surface receptors. For example, the NTD of the SARS-CoV-2 Spike interacts with the tyrosine-protein kinase receptor UFO (AXL) which appears to function as a co-receptor for human cell entry (Wang *et al*. 2021). The S2 subunit contains a heptad repeat region (including subregions HR1 and HR2) which mediate the fusion of the virion envelope with the host cell membrane during viral entry (Liu *et al*. 2004; Cui, Li and Shi 2019).

Being responsible for receptor binding and cellular entry, the evolution of the S gene is therefore key to host adaptation. It may be beneficial for coronaviruses to exchange either entire S genes, S1 subunit encoding portions of S genes, or smaller subdomains within the N-terminal domains of S1 during recombination, both because Spike is the main target of neutralising antibodies (Ou *et al*. 2020) and because the S gene is the main determinant of host species and host cell-type specificity (Lu, Wang and Gao 2015). Although recombination frequently transfers the entire S1 encoding region of the gene it is not uncommon in particular groups of viruses for it to transfer smaller subsections of the S1 (as can be seen with the red diagonals associated with the S genes of *Pedacoviruses* and *Merbecoviruses* in Figure2). In the *Alphacoronaviruses*, for example, recombination has involved transfer of the 5’ half of the NTD of S1 from transmissible gastroenteritis virus into canine coronavirus (type CCoV2b) (Decaro *et al*. 2009; Licitra, Duhamel and Whittaker 2014).

The S gene is also the only gene in which recombination cold-spots were detected in our breakpoint distribution analyses. Most noteworthy is that the 3’ 500 nucleotides of the S gene is the site of a conserved cold-spot detected in the *Igacoviruses, Pedacoviruses* and *Sarbecoviruses*. In the *Igacoviruses*, the coronavirus group with the richest full genome dataset in terms of both numbers of analysed sequences and their diversity, and within which the highest numbers of recombination breakpoints were detected (n=1703), our power to detect recombination cold-spots was greatest. Accordingly, recombination cold-spots were additionally detectable in the region of the S gene encoding the receptor-binding domain and dispersed throughout the 3’ half of the gene encoding the S2 subunit.

This arrangement of recombination cold-spots suggests that either basal recombination rates are suppressed within the NTD and S2 encoding regions of the S gene, or that recombination breakpoints falling within these regions tend to yield S genes that encode defective chimaeric Spike proteins. The NTD encoding region of the S gene is among the most genetically variable regions of coronavirus genomes and this alone might explain the relative absence of recombination breakpoints near the 5’ end of the S genes of *Igacoviruses, Nobecoviruses, Pedacoviruses*, and *Sarbecoviruses* (Archer *et al*. 2008; Boni *et al*. 2020). Similarly, the S2 encoding region of the S gene also tends to be more variable than most other coronavirus genome regions. However, the S2 subunit of Spike also contains multiple coevolved intra-protein amino acid interactions that are crucial for the cell-fusion functions of Spike (Bosch *et al*. 2003; Tang *et al*. 2020). It is also plausible, therefore, that the relative absence of detectable recombination breakpoints in the 3’ half of the S gene might be because recombinants carrying breakpoints falling within this region commonly express defective Spike proteins. In this regard, the S2 encoding region of the S gene may be a functional module that, while tending to retain its functionality when transferred by recombination as a complete unit into divergent genomic backgrounds (Wege *et al*. 1998), might be highly sensitive to recombination-induced disruptions of co-evolved amino acid interactions within S2 whenever recombination breakpoints fall within its boundaries.

### Selection likely disfavours recombinants expressing Spike proteins with disrupted folds

We used the SCHEMA method (Voigt *et al*. 2002; Lefeuvre *et al*. 2007) to more directly test for evidence of the inferred coronavirus recombination breakpoint distributions in the S gene having been impacted by natural selection disfavoring the survival of recombinants that express chimeric Spike proteins with disrupted folds. The only coronavirus proteins for which high resolution atomic coordinate data were available, and for which sufficient recombination breakpoint numbers were detected within their associated genome sites to perform the SCHEMA folding disruption test, were those of sequences in the *Merbecovirus, Sarbecovirus, Pedacovirus* and *Igacovirus* datasets.

We found that in the *Igacoviruses* and *Sarbecoviruses*, potential amino acid interactions within the Spike proteins expressed by observed recombinants have significantly fewer predicted structural impacts than would be expected under random recombination (p < 0.05; SCHEMA permutation test). It is noteworthy that the test result for the *Pedacoviruses* also approached significance (p = 0.079) but that for the *Merbecoviruses* displayed no such tendencies (p = 0.875; although it should be noted that, of the four datasets tested, this dataset had the lowest number of detected breakpoints in the S gene). This implies that, as has been suggested previously with *in vitro* recombination experiments involving the *Embecovirus*, murine coronavirus (Banner and Lai 1991), the *Igacoviruses* and *Sarbecoviruses* (and possibly also the *Pedacoviruses*) display lower degrees of predicted recombination-induced protein folding disruption in their expressed Spike proteins than would be expected under random recombination in the absence of selection. It should be stressed that our power to detect such “avoidance of protein folding disruption” signals was restricted to Spike and that it remains plausible that, given enough additional sequence data and more extensive atomic-resolution 3D structure information for other coronavirus proteins, many of these proteins might also display such signals.

### Indirect evidence that selection against protein misfolding impacts observable breakpoint distributions throughout coronavirus genomes

It would be expected that if natural selection tended to disfavour recombinants with misfolded proteins then breakpoints would tend to be found more frequently per non-coding nucleotide site than per amino acid encoding nucleotide site (Drummond *et al*. 2005). Also, it might be expected that, of the recombination breakpoints falling at amino acid encoding sites within genes, those falling at the edges of genes (for example in the first and last 5% of the coding sequence of a particular protein) might be less disruptive of coevolved intra-protein amino acid contacts that were crucial for correct folding than breakpoints falling within the middle regions of genes (Lefeuvre *et al*. 2007). If selection against misfolded proteins was impacting the distributions of recombination breakpoints throughout coronavirus genomes we would therefore expect that observed breakpoints might tend to fall more commonly: (1) in non-coding regions than in coding regions, and (2) at the edges of genes than in the middle parts of genes.

Accordingly, we found that the intergenic regions of the *Pedacoviruses, Embecoviruses, Nobecoviruses* and *Sarbecoviruses* all had significantly higher breakpoint densities (p < 0.05; permutation test; Table 1) than those in the protein-coding regions. Similarly, we detected that in the *Pedacoviruses, Embecoviruses, Sarbecoviruses* and *Igacoviruses*, detectable breakpoint densities were significantly higher in the beginning and ending 5% of coding regions than in the middle 90% of these regions (p < 0.05; permutation test; Table 2) with marginal significance observed in *Nobecoviruses* (p = 0.054; permutation test; Table 2).

**Table 1.** Comparison of detectable breakpoint numbers in non-coding regions and coding regions with rows in bold indicating subgenera with significantly more breakpoints in non-coding regions than would be expected under random recombination

| Subgenus | BPs* in non-coding regions | BPs in coding regions | Permutation p-val |
| --- | --- | --- | --- |
| ***Pedacovirus*** | **32** | **392** | **<0.001** |
| ***Embecovirus*** | **11** | **73** | **<0.001** |
| *Merbecovirus* | 1 | 66 | 0.660 |
| ***Nobecovirus*** | **4** | **79** | **0.012** |
| ***Sarbecovirus*** | **7** | **307** | **<0.001** |
| *Igacovirus* | 30 | 1683 | 0.650 |

\* BPs = Breakpoints.

**Table 2.**
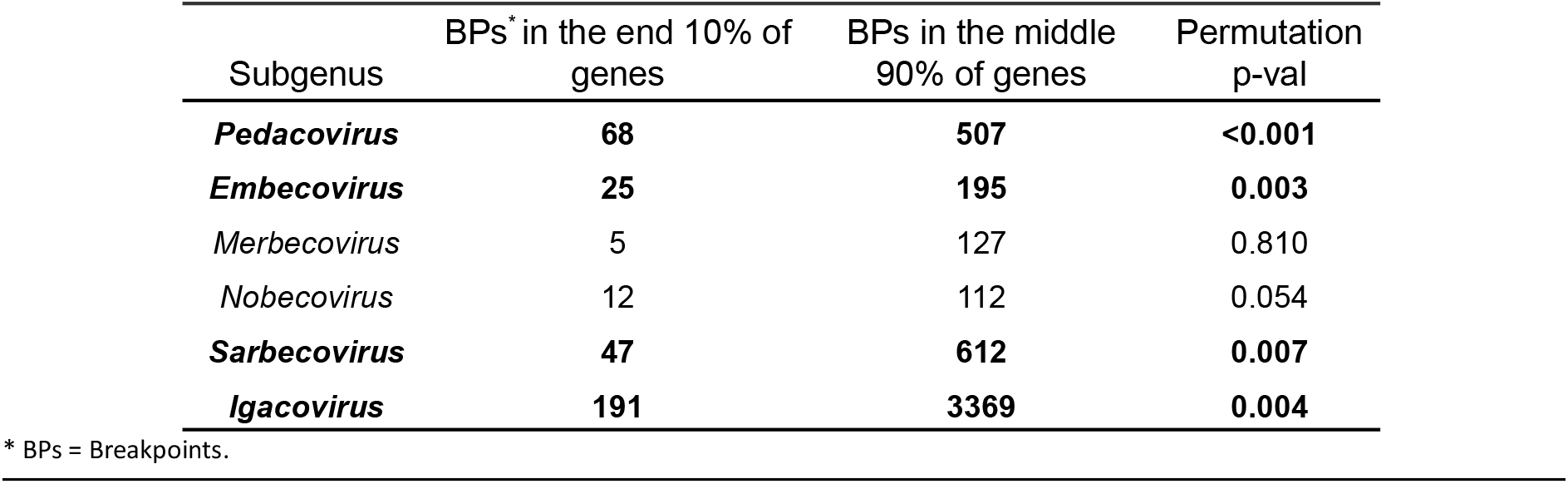
Breakpoint densities falling in the end 10% (5% each end) of genes vs the middle 90% of genes with rows in bold indicating subgenera with significantly higher numbers of detectable breakpoints in the ending 10% of genes than would be expected under random recombination

| Subgenus | BPs* in the end 10% of genes | BPs in the middle 90% of genes | Permutation p-val |
| --- | --- | --- | --- |
| ***Pedacovirus*** | **68** | **507** | **<0.001** |
| ***Embecovirus*** | **25** | **195** | **0.003** |
| *Merbecovirus* | 5 | 127 | 0.810 |
| *Nobecovirus* | 12 | 112 | 0.054 |
| ***Sarbecovirus*** | **47** | **612** | **0.007** |
| ***Igacovirus*** | **191** | **3369** | **0.004** |

\* BPs = Breakpoints.

Taken together the lower densities of breakpoints both within genes than in intergenic regions, and within the middle parts of genes than in the ends of genes is reminiscent of similar breakpoint distribution patterns detected in HIV (Simon-Loriere *et al*. 2010) and the members of various single-stranded DNA virus families (Lefeuvre *et al*. 2009) and is consistent with the hypothesis that in coronaviruses natural selection generally disfavours the survival of recombinants that express chimeric proteins with disrupted folds.

### ORF1a genome regions generally have lower breakpoint densities than other coding regions

There was a significantly lower density of breakpoints detected in ORF1a than in other coding regions of the genome for all six of the analysed subgenera (p < 0.05; permutation test; Table 3). The relatively low numbers of recombination events involving transfers of sequence fragments within this region is most notable in three of the *Betacoronaviruses* subgenera: *Embecovirus, Nobecovirus* and *Sarbecoviruses* (note the blue/cyan/green triangles associated with most of ORF1ab in these subgenera in Figure 2). Our results here are therefore consistent with previous observations that there is a significant tendency for recombination breakpoints to fall outside ORF1a in the human-infecting coronaviruses OC43 (an *Embecovirus*) and NL63 (an *Alphacoronavirus* in the subgenus *Setracovirus*) (Pollett *et al*. 2021).

**Table 3.** Individual genes and sub-gene regions with significantly lower numbers of detectable breakpoints than would be expected under random recombination

| Subgenus | Genome region | BPs* inside region | BPs outside region | permutation p-val |
| --- | --- | --- | --- | --- |
| *Pedacovirus* | ORF1a | 114 | 278 | 0.001 |
| *Embecovirus* | ORF1a | 43 | 130 | 0.001 |
| *Merbecovirus* | ORF1a | 43 | 130 | 0.001 |
| *Nobecovirus* | ORF1a | 14 | 65 | <0.001 |
| *Sarbecovirus* | ORF1a | 94 | 213 | <0.001 |
| *Igacovirus* | ORF1a | 667 | 1016 | 0.024 |
| *Nobecovirus* | plpro (nsp3) | 7 | 72 | 0.039 |
| *Sarbecovirus* | plpro (nsp3) | 49 | 258 | 0.031 |
| *Igacovirus* | plpro (nsp3) | 282 | 1401 | 0.016 |
| *Merbecovirus* | nsp4 | 0 | 66 | 0.035 |
| *Igacovirus* | nsp4 | 78 | 1605 | 0.002 |

\* BPs = Breakpoints.

Within ORF1a the regions encoding the nonstructural proteins (nsps) nsp3 (a papain-like cysteine protease), and nsp4 have particularly low densities of identified breakpoints in multiple different subgenera (*Nobecovirus, Sarbecovirus* and *Igacovirus* for nsp3 and *Merbecoviruses* and *Igacoviruses* for nsp4). Together with nsp6, nsp3 and nsp4 cooperatively modify the endoplasmic reticulum (ER) of coronavirus infected cells into vesicles with double membranes to which viral replication complexes are tethered (Knoops *et al*. 2008; Hartenian *et al*. 2020; Klein *et al*. 2020; Mohan and Wollert 2021).

It is plausible that the relatively low numbers of recombination events detectable in ORF1a are attributable to the high degree to which these components interact with one another (Stark *et al*. 2006; Li *et al*. 2021). It is expected that these interactions might rely on coevolved interaction motifs and that these proteins might therefore not function optimally if transferred into a genomic background within which they did not coevolve (Jain, Rivera and Lake 1999; Martin *et al*. 2005b).

### Breakpoints tend to fall at sites with lower than average GC content

To further our understanding of why, irrespective of selection, some coronavirus genomic sites might be more mechanistically predisposed to recombination than others, we tested breakpoint positions detected in each of the six analysed coronavirus datasets for associations with local GC contents (i.e. calculated proportions of all nucleotide residues that were G or C between 10 or 20 nucleotide sites up and downstream of detected breakpoint locations). High GC content is expected to potentially impact the frequencies at which recombination breakpoints occur in various ways such as (1) predisposing genome regions to form stable secondary structures that could cause pausing of RNA-Dependent RNA polymerase (RdRP) (Stark *et al*. 2006) (Experimental Evidence Codes | BioGRID 2021), (2) increasing the energy needed to break base-pairs during replication, and increasing the amount of time taken for RdRP to traverse these regions (Petes and Merker 2002; Sershen *et al*. 2011) and, if RdRPs disengages during replication, (3) increasing the probability of re-engagement through annealing with the same or a different template molecule (Lai 1990).

Contrary to expectations, but consistent with a recent report on recombination in coronaviruses (Pollett *et al*. 2021), we found that GC content within 20 nucleotides of breakpoint positions (Table 4) tended to be lower than expected under random recombination in all of the coronavirus datasets: significantly so in the *Sarbecovirus* and *Pedacovirus* datasets (P < 0.05; permutation test). When we repeated the test only considering GC contents within 10 nucleotides of recombination breakpoints (20 nt window in Table 4), the significant associations between breakpoint positions and lower GC content in *Sarbecoviruses* and *Pedacoviruses* were strengthened, and additionally, marginally significant associations with lower GC contents (0.05 < p < 0.1; permutation test) were detected in *Merbecoviruses, Embecoviruses* and *Igacoviruses*.

**Table 4.**
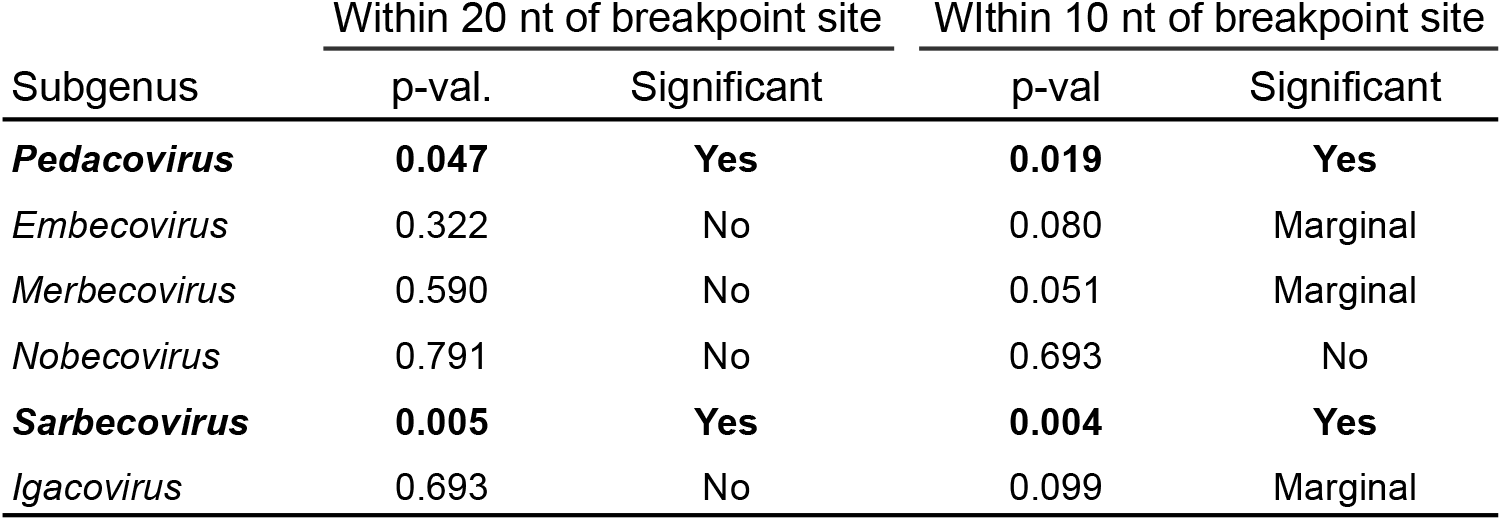
Associations between decreased GC content and detected recombination breakpoint sites with rows in bold indicating subgenera displaying average GC contents in the vicinity of breakpoint sites that are significantly lower than what would be expected under random recombination

| Subgenus | Within 20 nt of breakpoint site |  | Within 10 nt of breakpoint site |  |
| --- | --- | --- | --- | --- |
|  | p-val. | Significant | p-val | Significant |
| ***Pedacovirus*** | **0.047** | **Yes** | **0.019** | **Yes** |
| *Embecovirus* | 0.322 | No | 0.080 | Marginal |
| *Merbecovirus* | 0.590 | No | 0.051 | Marginal |
| *Nobecovirus* | 0.791 | No | 0.693 | No |
| ***Sarbecovirus*** | **0.005** | **Yes** | **0.004** | **Yes** |
| *Igacovirus* | 0.693 | No | 0.099 | Marginal |

### It is unclear whether sequence similarity directly influences the locations of recombination breakpoints

It has been previously found in other viruses that recombination breakpoint sites tend to occur more commonly at genome sites with elevated degrees of sequence conservation (van Vugt *et al*. 2001;Dazza *et al*. 2005; Archer *et al*. 2008). We therefore tested whether this pattern held for the six analyzed coronavirus subgenera.

Although recombination breakpoints in the *Sarbecovirus* and *Igacoviruses* datasets displayed a significant tendency to occur in genome regions displaying elevated degrees of average pairwise similarity among the analysed sequences (p < 0.007; permutation test; Table 5), for the *Pedacoviruses* and *Embecoviruses* the opposite was the case. In these subgenera, detectable recombination breakpoints have tended to fall in genome regions with lower degrees of average pairwise sequence similarity (p < 0.005; permutation test; Table 5). It is therefore unclear from our test whether pairwise sequence similarity within 10 or 20 nucleotides of prospective recombination breakpoint sites is a direct determinant of where breakpoints occur within coronavirus genomes.

**Table 5.** Association of breakpoint locations with higher/lower degrees of average pairwise sequence similarity with rows in bold indicating significant associations

| Subgenus | Within 20nt of breakpoint site |  | Within 10nt of breakpoint site |  |
| --- | --- | --- | --- | --- |
|  | Association with higher/lower similarity | p-val | Association with higher/lower similarity | p-val |
| ***Pedacovirus*** | **Lower** | **<0.001** | **Lower** | **<0.001** |
| ***Embecovirus*** | **Lower** | **0.006** | **Lower** | **0.007** |
| *Merbecovirus* | Higher | 0.184 | Higher | 0.192 |
| *Nobecovirus* | Higher | 0.465 | Higher | 0.475 |
| ***Sarbecovirus*** | **Higher** | **0.005** | **Higher** | **0.005** |
| ***Igacovirus*** | **Higher** | **<0.001** | **Higher** | **<0.001** |

It is noteworthy in this regard that there are substantial variations in degrees of sequence conservation across the analysed sequence datasets with, for example, the genome regions corresponding to the recombination breakpoint hot-spot immediately upstream of the S gene in the *Nobecovirus, Sarbecovirus* and *Igacovirus* datasets (all with a tendency for breakpoints to fall at more conserved sites) displaying among the highest degrees of sequence conservation within these datasets (Figure 3). Conversely, for the *Pedacoviruses* and *Embecoviruses* datasets (both with a tendency for breakpoints to fall at less conserved sites) the corresponding recombination hot-spots upstream of the S gene start codon fall at genome sites that have among the lowest degrees of genome-wide conservation in these datasets (Figure 3). It is therefore likely that, for this conserved hot-spot at least, sequence similarity has not been a primary determinant of where breakpoints have occurred.

**Figure 3.**
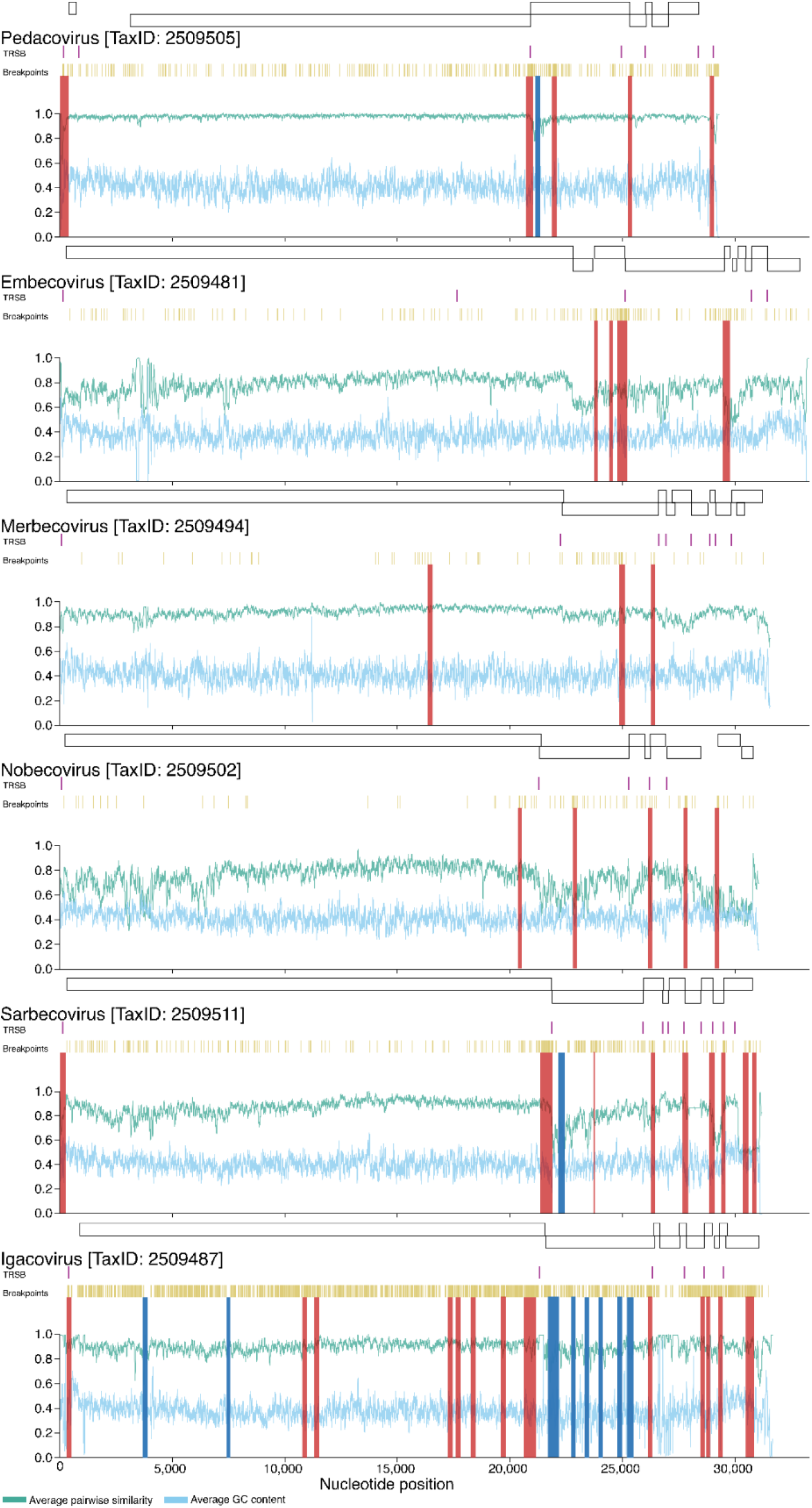
Regional variations in average pairwise sequence similarity (green) and GC content (blue) across coronavirus genomes. The plotted values indicate the pairwise sequence similarities and GC proportions within a moving 40 nucleotide window. Also indicated are the locations of the main genes (above each graph), transcriptional regulatory sequences (TRSs; in purple), identified breakpoint locations (in mustard), potential recombination hotspots (in red) and potential recombination cold-spots (in blue).

### There is a strong association between recombination breakpoint locations and those of transcriptional regulatory sequences

Coronavirus transcription involves template switching at specific genome sites, called transcriptional regulatory sequences (TRSs) (Yang *et al*. 2021), previously called the intergenic sequence (Alonso *et al*. 2002). A possible link between template switching during gene expression and the genomic sites where recombination breakpoints occur during genome replication has been noted previously for coronaviruses in general (Zúñiga *et al*. 2004; Sola *et al*. 2015) and SARS-CoV specifically (Graham *et al*. 2018). Template switching is prone to occur during transcription of coronavirus negative genome strands whenever RdRp encounters the TRS sequences that are commonly found upstream of various genes. Because these “body TRS” (or TRS-B;) (Alonso *et al*. 2002; Sola *et al*. 2015) sites are involved in frequent template switching during transcription, it has been suggested that these sites might also promote template switching during genome replication (Graham et al. 2018) and, therefore, that they might colocalize with recombination hot-spots (Yang *et al*. 2021).

We used the SuPER method (Yang *et al*. 2021) to detect potential TRS-B sites in each of our six coronavirus datasets. Whereas SuPER can use RNA-seq data to precisely locate TRS-B sites, in our case we used previously identified TRS-L sequences (Yang *et al*. 2021) to find and annotate likely TRS-B sites within the six analysed coronavirus datasets.

We found strong evidence for associations between the locations of conserved TRS-B sites (i.e. those detected in >75% of the analysed sequences in each dataset) and the locations of detected recombination breakpoints in the *Pedacoviruses, Igacoviruses, Embecoviruses* and *Sarbecoviruses* (p < 0.05; permutation test; Table 6). These associations were detectable when we varied the required proximity between breakpoints and potential TRS-B sites to be considered a match from between 2 and 46 nucleotides.

**Table 6.** Associations between transcription regulatory sequence (TRS) sites and the locations of detected recombination breakpoints with p-values in bold indicating significant associations of TRS sites with higher breakpoint numbers

| Subgenus | Within 46 nts<br>p-val | Within 21 nts<br>p-val | Within 9 nts<br>p-val | Within 2 nts<br>p-val |
| --- | --- | --- | --- | --- |
| *Pedacovirus* | **<0.001** | **<0.001** | **<0.001** | **<0.001** |
| *Embecovirus* | **<0.001** | **<0.001** | **<0.001** | **<0.001** |
| *Merbecovirus* | 0.117 | 0.178 | 0.210 | 0.806 |
| *Nobecovirus* | **0.014** | **0.039** | **0.020** | 0.478 |
| *Sarbecovirus* | **<0.001** | **<0.001** | **<0.001** | **0.005** |
| *Igacovirus* | **<0.001** | **<0.001** | **<0.001** | **<0.001** |

Given the large detectable recombination breakpoint hot-spots directly upstream of the S gene in most of the analysed subgenera datasets and the TRS-B sequences that map near these hot-spots, it was possible that the associations detected between TRS locations and breakpoint positions could have been attributable entirely to the TRS-B sites upstream of the spike gene. To determine if this was the case, we repeated the association test (25nt window size) but this time with the TRS upstream of Spike removed from the analysis. We observed a minimal decrease in the significance of the association between TRS-B sites and recombination breakpoint positions, indicating that the initial result was not simply being driven by the coincidental colocalization of the S gene associated TRS-B site and the conserved recombination hot-spot upstream of the S gene in most of the analysed datasets.

We reran the TRS-B association tests with the positions of the TRS-B sites randomly shifted along the genome. The script takes as input the alignment file for each of the six datasets and places five to ten “false” TRS-B sites across each genome (the exact number corresponding for each subgenus dataset to the “true” TRS-B number for that dataset). When considering breakpoint probability distributions and an analysis window of 25 nucleotides, there was a significant absence of breakpoints within 12 nucleotides of TRS-B sites in the *Embecovirus* and *Sarbecovirus* datasets and neither significantly more or less breakpoints in close proximity to TRS-B sites in any of the other datasets. Both these results, along with our previous tests, are strong evidence that recombination breakpoints in coronaviruses generally tend to cluster at TRS-B sites.

However, given we have found that detectable recombination breakpoints tend to fall near the edges of genes in the same four subgenera in which we detected an association between TRS-B locations and recombination breakpoints (*Pedacovirus, Embecovirus, Sarbecovirus* and *Igacovirus*), this association between breakpoint locations and TRS-B sites might simply be attributable to the fact that TRS-B sites also tend to fall at the edges of genes. We therefore attempted to determine whether the association between breakpoint locations and TRS-B sites were still evident if we controlled for the colocalization of these sites at the edges of genes. We were specifically interested in whether the presence/absence of a TRS-B site immediately upstream of a gene was associated with the presence/absence of a recombination breakpoint hot-spot upstream of the gene. Considering only the TRS-B sites and recombination breakpoint hot-spots falling either in intergenic regions or within 300 nucleotides of the beginning of genes we found a significant association between the presence of a TRS-B site near the beginning of a gene and the presence of a hot-spot near that location (p = 0.0392, Chi-square test with N-1 correction). Therefore suggesting that, for the *Pedacovirus, Sarbecovirus, Igacovirus* and *Embecovirus* datasets at least, the significant association we found between TRS-B sites and recombination breakpoint locations was not merely attributable to a coincidental tendency for breakpoints and TRS-B sites to colocalize near the edges of genes.

## Conclusion

Across all of the tests that we performed, viruses in the different analysed coronavirus genera displayed similar patterns of recombination (Table 7). The most strikingly similar of these patterns were those observed in the *Sarbecoviruses* (members of the *Betacoronavirus* genus) and the *Pedacoviruses* (members of the Alphacoronavirus genus). These mostly concordant patterns indicate that the processes that yield and select recombinant coronaviruses are likely broadly conserved across the three analysed coronavirus genera.

**Table 7.** Conserved patterns of recombination across various coronavirus subgenera. Rows each contain the result of either a statistical test or the presence/absence of a particular characteristic of recombination (such as the presence of a hot-spot at a specific genome location): BP = breakpoint; blue = significant association or presence of characteristic; light blue = marginally significant association; pink = no significant association or absence of characteristic; white = untested.

|  | *Pedacovirus* | *Sarbecovirus* | *Nobecovirus* | *Embecovirus* | *Igacovirus* | *Merbecovirus* |
| --- | --- | --- | --- | --- | --- | --- |
| Hot-spot at 5' end of genome | blue | blue | blue | pink | blue | pink |
| Hot-spot upstream of S gene | blue | blue | blue | blue | blue | pink |
| Hot-spot in 5' half of S gene | blue | blue | blue | pink | pink | blue |
| Cold-spot at 5' end of S gene | blue | blue | pink | pink | blue | pink |
| More BPs in intergenic regions | blue | blue | blue | blue | pink | pink |
| Fewer BPs in middle of genes | blue | blue | light blue | blue | blue | pink |
| Higher BP density in lower GC regions | blue | blue | pink | light blue | pink | light blue |
| Higher BP density near TRS-B sites | blue | blue | light blue | blue | blue | pink |
| Avoidance of S gene fold disruption | light blue | blue | white | white | blue | pink |

The subgenus dataset displaying the least concordant recombination patterns was that of the *Merbecoviruses*. It is unclear to us why the *Mercbecovirus* dataset displays recombination breakpoint patterns that differ from the other analysed datasets: it is not an outlier among the datasets in terms of the numbers of sequences analysed or the average pairwise similarities of these sequences, but the dataset does have the lowest number of detectable recombination events. It is therefore possible that either the processes that generate recombinant genomes, the genetic factors that determine the viability of recombinants, or the epidemiological and evolutionary processes that impact the survival of recombinants, might differ somewhat between the *Merbecoviruses* and most other coronaviruses.

Nevertheless, the non-random and mostly conserved recombination patterns that we and others have detected in various coronavirus subgenera are likely shaped both by evolutionarily conserved variations in the mechanistic predispositions of different genome regions to recombination and by shared selective processes disfavoring the survival of recombinants that express improperly folded proteins. There are two non-exclusive explanations for why coronavirus genome sites that are mechanistically predisposed to recombination (such as those of TRS-B sequences) tend to coincide with sites where recombination seems to have had a minimal impact on protein folding: (1) negative selection over the short-term may be so efficient at purging all viral variants with recombination-induced protein misfolding that such variants are only rarely sequenced; and/or (2) longer-term selection, possibly acting since the most recent common ancestor of all known coronaviruses, may have yielded coronavirus genomes that are configured such that they are mechanistically predisposed to only recombine at sites where recombination breakpoints are minimally disruptive of protein folding. When high-resolution maps of amino acid contacts within coronavirus protein complexes become available, and when the conserved nucleotide interactions within biologically functional RNA structural elements in a diverse enough array of coronavirus genomes have been identified, it should also be possible to determine the degree to which selection acting over the short- and/or long-terms to preserve these other categories of coevolved intra-genome interactions have impacted observable coronavirus recombination patterns.

## Supporting information

RDP5_Supplementary_Data

## Acknowledgements

ADK and PS were supported by a University of Cape Town Masters Research Scholarship [960000000760].

RL was supported by the South African National Research Foundation.

DPM was supported by the Wellcome Trust [222574/Z/21/Z].

SLKP was supported by the U.S. National Institutes of Health [R01 AI134384 and AI140970] and the US National Science Foundation [RAPID 2027196 NSF/DBI,BIO].

SL was supported by the Medical Research Council of the United Kingdom [MC_UU_12014/12].

OAM was supported by the Wellcome Trust [206369/Z/17/Z].

JDZ was supported by the U.S. National Institutes of Health [R01 AI134384 and AI140970].

MZ, IA, DR, VK, MJS, GH and BM were not supported.

## Data available in Supplementary Data

## Supplementary Data

**Supplementary Table 1.** Coronavirus whole genome nucleotide sequence dataset characteristics

| Subgenus | Genus | Common animal hosts | Number of sequences | Mean pairwise similarity |
| --- | --- | --- | --- | --- |
| *Pedacovirus* | *Alphacoronavirus* | Bat, Porcine | 412 | 98% |
| *Merbecovirus* | *Betacoronavirus* | Bat, Camel, Human | 191 | 92% |
| *Nobecovirus* | *Betacoronavirus* | Bat | 16 | 75% |
| *Sarbecovirus* | *Betacoronavirus* | Bat, Pangolin, Human | 158 | 86% |
| *Embecovirus* | *Betacoronavirus* | Rodent, Bovine, Porcine, Rabbit, Deer, Human | 181 | 78% |
| *Igacovirus* | *Gammacoronavirus* | Pigeon, Fowl, Pheasant | 410 | 89% |

**Supplementary Table 2:** TRS-B association test

|  | Gene boundaries without TRS-Bs | hot-spots | Gene boundaries with TRS-Bs | hot-spots |
| --- | --- | --- | --- | --- |
| *Sarbecovirus* | 1 | 1 | 9 | 6 |
| *Pedacovirus* | 3 | 1 | 5 | 3 |
| *Igacovirus* | 6 | 0 | 6 | 5 |
| *Embecovirus* | 7 | 3 | 4 | 1 |
| total | 17 | 5 | 24 | 15 |
| N (boundary regions) | 41 |  |  |  |

**Supplementary Table 3:** Contingency table for TRS-Bs and hot-spots at gene boundaries

|  | w/o TRS-B | With TRS-B | Marginal Totals |
| --- | --- | --- | --- |
| No hot-spot | 12 | 9 | 21 |
| Hot-spot | 5 | 15 | 20 |
| Marginal Totals | 17 | 24 | 41 |

## References

Alonso S, Izeta A, Sola I et al. Transcription Regulatory Sequences and mRNA Expression Levels in the Coronavirus Transmissible Gastroenteritis Virus. J Virol 2002;76:1293–308.

Anthony SJ, Gilardi K, Menachery VD et al. Further Evidence for Bats as the Evolutionary Source of Middle East Respiratory Syndrome Coronavirus. Schultz-Cherry S (ed.). mBio 2017;8, DOI: 10.1128/mBio.00373-17.

Archer J, Pinney JW, Fan J et al. Identifying the Important HIV-1 Recombination Breakpoints. PLOS Comput Biol 2008;4:e1000178.

Banner LR, Lai MM. Random nature of coronavirus RNA recombination in the absence of selection pressure. Virology 1991;185:441–5.

Belouzard S, Millet JK, Licitra BN et al. Mechanisms of Coronavirus Cell Entry Mediated by the Viral Spike Protein. Viruses 2012;4:1011–33.

Berman H, Henrick K, Nakamura H. Announcing the worldwide Protein Data Bank. Nat Struct Biol 2003;10:980.

Boni MF, Lemey P, Jiang X et al. Evolutionary origins of the SARS-CoV-2 sarbecovirus lineage responsible for the COVID-19 pandemic. Nat Microbiol 2020 511 2020;5:1408–17.

Bosch BJ, van der Zee R, de Haan CAM et al. The Coronavirus Spike Protein Is a Class I Virus Fusion Protein: Structural and Functional Characterization of the Fusion Core Complex. J Virol 2003;77:8801–11.

Cheng C-P, Nagy PD. Mechanism of RNA Recombination in Carmo- and Tombusviruses: Evidence for Template Switching by the RNA-Dependent RNA Polymerase In Vitro. J Virol 2003;77:12033.

Coronaviridae - Positive Sense RNA Viruses - Positive Sense RNA Viruses (2011) - ICTV. ICTV 2011.

Crameri A, Raillard S, Bermudez E et al. DNA shuffling of a family of genes from diverse species accelerates directed evolution. Nature 1998;391:288–91.

Cui J, Li F, Shi Z-L. Origin and evolution of pathogenic coronaviruses. Nat Rev Microbiol 2019;17:181–92.

Dazza M-C, Ekwalanga M, Nende M et al. Characterization of a Novel *vpu* -Harboring Simian Immunodeficiency Virus from a Dent’s Mona Monkey (Cercopithecus mona denti). J Virol 2005;79:8560–71.

Decaro N, Mari V, Campolo M et al. Recombinant Canine Coronaviruses Related to Transmissible Gastroenteritis Virus of Swine Are Circulating in Dogs. J Virol 2009;83:1532–7.

Decaro N, Mari V, Elia G et al. Full-length genome analysis of canine coronavirus type I. Virus Res 2015;210:100–5.

Denison MR, Graham RL, Donaldson EF et al. Coronaviruses: An RNA proofreading machine regulates replication fidelity and diversity. RNA Biol 2011;8:270–9.

Drummond DA, Silberg JJ, Meyer MM et al. On the conservative nature of intragenic recombination. Proc Natl Acad Sci 2005;102:5380–5.

Dudas G, Rambaut A. MERS-CoV recombination: implications about the reservoir and potential for adaptation. Virus Evol 2016;2:vev023.

Emam M, Oweda M, Antunes A et al. Positive selection as a key player for SARS-CoV-2 pathogenicity: Insights into ORF1ab, S and E genes. Virus Res 2021;302:198472. Experimental Evidence Codes | BioGRID. BioGRID 2021.

Forni D, Cagliani R, Sironi M. Recombination and Positive Selection Differentially Shaped the Diversity of Betacoronavirus Subgenera. Viruses 2020;12:1313.

Gibbs MJ, Armstrong JS, Gibbs AJ. Sister-Scanning: a Monte Carlo procedure for assessing signals in recombinant sequences. Bioinformatics 2000;16:573–82.

Goldstein SA, Brown J, Pedersen BS et al. Extensive Recombination-Driven Coronavirus Diversification Expands the Pool of Potential Pandemic Pathogens., 2021:2021.02.03.429646.

Graham RL, Deming DJ, Deming ME et al. Evaluation of a recombination-resistant coronavirus as a broadly applicable, rapidly implementable vaccine platform. Commun Biol 2018;1:179.

Hartenian E, Nandakumar D, Lari A et al. The molecular virology of coronaviruses. J Biol Chem 2020;295:12910–34.

Hatcher EL, Zhdanov SA, Bao Y et al. Virus Variation Resource - improved response to emergent viral outbreaks. Nucleic Acids Res 2017;45:D482–90.

Heath L, Van Der Walt E, Varsani A et al. Recombination Patterns in Aphthoviruses Mirror Those Found in Other Picornaviruses. J Virol 2006;80:11827–32.

Herrewegh AAPM, Smeenk I, Horzinek MC et al. Feline Coronavirus Type II Strains 79-1683 and 79-1146 Originate from a Double Recombination between Feline Coronavirus Type I and Canine Coronavirus. J Virol 1998;72:4508–14.

Hussin JG, Hodgkinson A, Idaghdour Y et al. Recombination affects accumulation of damaging and disease-associated mutations in human populations. Nat Genet 2015 474 2015;47:400–4.

Jain R, Rivera MC, Lake JA. Horizontal gene transfer among genomes: The complexity hypothesis. Proc Natl Acad Sci 1999;96:3801–6.

Jaroszewski L, Iyer M, Alisoltani A et al. The interplay of SARS-CoV-2 evolution and constraints imposed by the structure and functionality of its proteins. Punta M (ed.). PLOS Comput Biol 2021;17:e1009147.

Jia W, Karaca K, Parrish CR et al. A novel variant of avian infectious bronchitis virus resulting from recombination among three different strains. Arch Virol 1995;140:259–71.

Katoh K, Standley DM. MAFFT Multiple Sequence Alignment Software Version 7: Improvements in Performance and Usability. Mol Biol Evol 2013;30:772.

Keck JG, Matsushima GK, Makino S et al. In vivo RNA-RNA recombination of coronavirus in mouse brain. J Virol 1988a;62:1810–3.

Keck JG, Soe LH, Makino S et al. RNA recombination of murine coronaviruses: recombination between fusion-positive mouse hepatitis virus A59 and fusion-negative mouse hepatitis virus 2. J Virol 1988b;62:1989–98.

Klein S, Cortese M, Winter SL et al. SARS-CoV-2 structure and replication characterized by in situ cryo-electron tomography. Nat Commun 2020;11:5885.

Knoops K, Kikkert M, Worm SHE van den et al. SARS-Coronavirus Replication Is Supported by a Reticulovesicular Network of Modified Endoplasmic Reticulum. PLOS Biol 2008;6:e226.

Krumm ZA, Lloyd GM, Francis CP et al. Precision therapeutic targets for COVID-19. Virol J 2021 181 2021;18:1–22.

Kusters JG, Jager EJ, Niesters HG et al. Sequence evidence for RNA recombination in field isolates of avian coronavirus infectious bronchitis virus. Vaccine 1990;8:605–8.

Lai MM. Coronavirus: organization, replication and expression of genome. Annu Rev Microbiol 1990;44:303–303.

Lai MMC. Recombination in large RNA viruses: Coronaviruses. 1996.

Lam H, Ratmann O, Boni M. Improved Algorithmic Complexity for the 3SEQ Recombination Detection Algorithm. Mol Biol Evol 2018;35:247–51.

Lednicky JA, Tagliamonte MS, White SK et al. Isolation of a Novel Recombinant Canine Coronavirus from a Visitor to Haiti: Further Evidence of Transmission of Coronaviruses of Zoonotic Origin to Humans. Clin Infect Dis 2021, DOI: 10.1093/cid/ciab924.

Lee CW, Jackwood MW. Evidence of genetic diversity generated by recombination among avian coronavirus IBV. Arch Virol 2000;145:2135–48.

Lefeuvre P, Lett J-M, Reynaud B et al. Avoidance of Protein Fold Disruption in Natural Virus Recombinants. PLOS Pathog 2007;3:e181.

Lefeuvre P, Lett J-M, Varsani A et al. Widely Conserved Recombination Patterns among Single-Stranded DNA Viruses. J Virol 2009;83:2697–707.

Li BX, Ge JW, Li YJ. Porcine aminopeptidase N is a functional receptor for the PEDV coronavirus. Virology 2007;365:166–72.

Li J, Guo M, Tian X et al. Virus-Host Interactome and Proteomic Survey Reveal Potential Virulence Factors Influencing SARS-CoV-2 Pathogenesis. Med 2021;2:99–112.e7.

Licitra B, Duhamel G, Whittaker G. Canine Enteric Coronaviruses: Emerging Viral Pathogens with Distinct Recombinant Spike Proteins. Viruses 2014;6:3363–76.

Liu P, Shi L, Zhang W et al. Prevalence and genetic diversity analysis of human coronaviruses among cross-border children. Virol J 2017 141 2017;14:1–8.

Liu S, Xiao G, Chen Y et al. Interaction between heptad repeat 1 and 2 regions in spike protein of SARS-associated coronavirus: implications for virus fusogenic mechanism and identification of fusion inhibitors. The Lancet 2004;363:938–47.

Lu G, Wang Q, Gao GF. Bat-to-human: spike features determining "host jump" of coronaviruses SARS-CoV, MERS-CoV, and beyond. Trends Microbiol 2015;23:468–78.

Lytras S, Hughes J, Martin D et al. Exploring the Natural Origins of SARS-CoV-2 in the Light of Recombination., 2021:2021.01.22.427830.

Makino S, Keck J, SA S et al. High-frequency RNA recombination of murine coronaviruses. J Virol 1986;57:729–37.

Manfredonia I, Nithin C, Ponce-Salvatierra A et al. Genome-wide mapping of SARS-CoV-2 RNA structures identifies therapeutically-relevant elements. Nucleic Acids Res 2020;48:12436–52.

Martin D, Posada D, Crandall KA et al. A Modified Bootscan Algorithm for Automated Identification of Recombinant Sequences and Recombination Breakpoints. https://home.liebertpub.com/aid 2005a;21:98–102.

Martin D, Rybicki E. RDP: detection of recombination amongst aligned sequences. Bioinformatics 2000;16:562–3.

Martin D, Varsani A, Roumagnac P et al. RDP5: a computer program for analyzing recombination in, and removing signals of recombination from, nucleotide sequence datasets. Virus Evol 2021;7:veaa087.

Martin D, van der Walt E, Posada D et al. The Evolutionary Value of Recombination Is Constrained by Genome Modularity. PLOS Genet 2005b;1:e51.

Maynard Smith J. Analyzing the mosaic structure of genes. J Mol Evol 1992 342 1992;34:126–9.

Meyer MM, Silberg JJ, Voigt CA et al. Library analysis of SCHEMA-guided protein recombination. Protein Sci 2003;12:1686–93.

Mohan J, Wollert T. Membrane remodeling by SARS-CoV-2 – double-enveloped viral replication. Fac Rev 2021;10:17.

Muller H. The relation of recombination to mutational advance. Mutat Res 1964;106:2–9.

Ou X, Liu Y, Lei X et al. Characterization of spike glycoprotein of SARS-CoV-2 on virus entry and its immune cross-reactivity with SARS-CoV. Nat Commun 2020;11:1620.

Petes TD, Merker JD. Context Dependence of Meiotic Recombination Hotspots in Yeast: The Relationship Between Recombination Activity of a Reporter Construct and Base Composition. Genetics 2002;162:2049–52.

Pettersen E, Goddard T, Huang C et al. UCSF Chimera–a visualization system for exploratory research and analysis. J Comput Chem 2004;25:1605–12.

Pollett S, Conte MA, Sanborn M et al. A comparative recombination analysis of human coronaviruses and implications for the SARS-CoV-2 pandemic. Sci Rep 2021;11:17365.

Raj VS, Mou H, Smits SL et al. Dipeptidyl peptidase 4 is a functional receptor for the emerging human coronavirus-EMC. Nature 2013;495:251–4.

Reusken CB, Raj VS, Koopmans MP et al. Cross host transmission in the emergence of MERS coronavirus. Curr Opin Virol 2016;16:55–62.

Rossum G van, Drake FL. The Python Language Reference. Release 3.0.1 [Repr.]. Hampton, NH: Python Software Foundation, 2010.

Sawyer S. Statistical tests for detecting gene conversion. Mol Biol Evol 1989;6:526–38.

Sershen CL, Mell JC, Madden SM et al. Superhelical Duplex Destabilization and the Recombination Position Effect. Lustig AJ (ed.). PLoS ONE 2011;6:e20798.

Siegfried NA, Busan S, Rice GM et al. RNA motif discovery by SHAPE and mutational profiling (SHAPE-MaP). Nat Methods 2014;11:959–65.

Simon-Loriere E, Galetto R, Hamoudi M et al. Molecular Mechanisms of Recombination Restriction in the Envelope Gene of the Human Immunodeficiency Virus. PLoS Pathog 2009;5:1000418.

Simon-Loriere E, Martin D, Weeks K et al. RNA structures facilitate recombination-mediated gene swapping in HIV-1. J Virol 2010;84:12675–82.

Sola I, Almazán F, Zúñiga S et al. Continuous and Discontinuous RNA Synthesis in Coronaviruses. Annu Rev Virol 2015;2:265–88.

Song S, Ma L, Zou D et al. The Global Landscape of SARS-CoV-2 Genomes, Variants, and Haplotypes in 2019nCoVR. Genomics Proteomics Bioinformatics 2021, DOI: 10.1016/J.GPB.2020.09.001.

Stark C, Breitkreutz B-J, Reguly T et al. BioGRID: a general repository for interaction datasets. Nucleic Acids Res 2006;34:D535–9.

Su S, Wong G, Shi W et al. Epidemiology, Genetic Recombination, and Pathogenesis of Coronaviruses. Trends Microbiol 2016;24:490–502.

Tang T, Bidon M, Jaimes JA et al. Coronavirus membrane fusion mechanism offers a potential target for antiviral development. Antiviral Res 2020;178:104792.

Vlasova AN, Diaz A, Damtie D et al. Novel Canine Coronavirus Isolated from a Hospitalized Patient With Pneumonia in East Malaysia. Clin Infect Dis 2021, DOI: 10.1093/cid/ciab456.

Voigt CA, Martinez C, Wang Z-G et al. Protein building blocks preserved by recombination. Nat Struct Biol 2002;9:553–8.

van Vugt JJFA, Storgaard T, Oleksiewicz MB et al. High frequency RNA recombination in porcine reproductive and respiratory syndrome virus occurs preferentially between parental sequences with high similarity. J Gen Virol 2001;82:2615–20.

Wan Y, Shang J, Graham R et al. Receptor Recognition by the Novel Coronavirus from Wuhan: an Analysis Based on Decade-Long Structural Studies of SARS Coronavirus. Gallagher T (ed.). J Virol 2020;94, DOI: 10.1128/JVI.00127-20.

Wang L, Junker D, Collisson EW. Evidence of natural recombination within the S1 gene of infectious bronchitis virus. Virology 1993;192:710–6.

Wang S, Qiu Z, Hou Y et al. AXL is a candidate receptor for SARS-CoV-2 that promotes infection of pulmonary and bronchial epithelial cells. Cell Res 2021;31:126–40.

Wang W, Lin X-D, Guo W-P et al. Discovery, diversity and evolution of novel coronaviruses sampled from rodents in China. Virology 2015;474:19–27.

Wang W, Lin X-D, Zhang H-L et al. Extensive genetic diversity and host range of rodent-borne coronaviruses. Virus Evol 2020;6:veaa078.

Wege H, Stühler A, Lassmann H et al. Coronavirus infection and demyelination: Sequence conservation of the S-gene during persistent infection of Lewis-rats. Adv Exp Med Biol 1998;440:767–73.

Wesley RD. The S gene of canine coronavirus, strain UCD-1, is more closely related to the S gene of transmissible gastroenteritis virus than to that of feline infectious peritonitis virus. Virus Res 1999;61:145–52.

White JM, Delos SE, Brecher M et al. Structures and Mechanisms of Viral Membrane Fusion Proteins: Multiple Variations on a Common Theme. Crit Rev Biochem Mol Biol 2008;43:189–219.

Woo PCY, Huang Y, Lau SKP et al. Coronavirus Genomics and Bioinformatics Analysis. Viruses 2010;2:1804–20.

Wrapp D, Wang N, Corbett KS et al. Cryo-EM structure of the 2019-nCoV spike in the prefusion conformation. Science 2020;367:1260–3.

Xia S, Liu M, Wang C et al. Inhibition of SARS-CoV-2 (previously 2019-nCoV) infection by a highly potent pan-coronavirus fusion inhibitor targeting its spike protein that harbors a high capacity to mediate membrane fusion. Cell Res 2020;30:343–55.

Yang D, Leibowitz JL. The structure and functions of coronavirus genomic 3' and 5' ends. Virus Res 2015;206:120–33.

Yang Y, Yan W, Hall AB et al. Characterizing Transcriptional Regulatory Sequences in Coronaviruses and Their Role in Recombination. Mol Biol Evol 2021;38:1241–8.

Yeager CL, Ashmun RA, Williams RK et al. Human aminopeptidase N is a receptor for human coronavirus 229E. Nature 1992;357:420–2.

Zehr JD, Kosakovsky Pond SL, Martin DP et al. Recent Zoonotic Spillover and Tropism Shift of a Canine Coronavirus Is Associated with Relaxed Selection and Putative Loss of Function in NTD Subdomain of Spike Protein. Evolutionary Biology, 2021.

Zhu Z, Meng K, Liu G et al. A database resource and online analysis tools for coronaviruses on a historical and global scale. Database 2021;2020, DOI: 10.1093/DATABASE/BAAA070.

Zúñiga S, Sola I, Alonso S et al. Sequence Motifs Involved in the Regulation of Discontinuous Coronavirus Subgenomic RNA Synthesis. J Virol 2004;78:980–94.

